# Using a ‘one strain-many compounds’ (OSMAC) approach to screen a collection of diverse fungi from Aotearoa New Zealand for antibacterial activity against *Escherichia coli*

**DOI:** 10.1101/2025.05.05.652310

**Authors:** Shara van de Pas, Melissa M. Cadelis, Alexander B.J. Grey, Jessica M. Flemming, Duckchul Park, Thomas Lumley, Bevan S. Weir, Brent R. Copp, Siouxsie Wiles

## Abstract

There is an urgent need to identify new chemical compounds with novel modes of action to help manage the antimicrobial resistance crisis. Fungi are prolific producers of secondary metabolites, including those with antimicrobial properties, and contain biosynthetic gene clusters that awaken only under certain growth conditions. In recent years, a wealth of novel fungal biosynthetic pathways and compounds have been identified, suggesting fungi remain a viable source for developing new antimicrobials. The International Collection of Microorganisms from Plants (ICMP) contains thousands of fungi and bacteria primarily sourced from Aotearoa New Zealand. Here, we report the results of our efforts to screen 32 fungal ICMP isolates for activity against *Escherichia coli*, a leading cause of deaths attributable to antimicrobial resistance. We used a ‘one strain-many compounds’ (OSMAC) approach, growing the ICMP isolates on seven different media with different pH and various carbon and nitrogen sources. We also tested the isolates for activity at various ages. Our results indicate that several of the tested fungi possess anti-*E. coli* activity and are suitable for further study, including the aero-aquatic Ascomycete *Hyaloscypha* sp. ICMP 16864. Our results also provide further strong evidence for the impact of media on both fungal growth and bioactivity.

## Introduction

It is 80 years since Sir Alexander Fleming received the Nobel Prize in Physiology or Medicine for discovering the antibiotic penicillin. Penicillin and other antibiotics have become crucial medical tools in the intervening years. As well as being given to treat those with a bacterial infection, antibiotics are also used to prevent infection in vulnerable patients, including people undergoing chemotherapy or surgery. In his Nobel Lecture in December 1945, Fleming warned that it was “not difficult to make microbes resistant to penicillin” and advised against using the drug negligently [1]. Since then, resistance to antibiotics and other antimicrobials has become a major threat to human health around the world [2], exacerbated even further by the coronavirus disease 2019 (COVID-19) pandemic [3–5].

Before the COVID-19 pandemic, an assessment of the global burden of antimicrobial resistance estimated that in 2019, almost 5 million deaths were associated with bacterial resistance to antibiotics [6]. This figure includes 1.27 million deaths directly attributable to resistance, with the leading cause being *Escherichia coli* [6]. While the share of deaths caused by *E. coli* varied by region, in the high-income super-region, this bacterium alone was linked to almost one in four deaths attributable to antimicrobial resistance [6].

To help manage this crisis, there is an urgent need to identify new chemical compounds with novel modes of antimicrobial action [7, 8]. Most antibiotics in the clinic today derive from compounds identified from soil microbes, beginning with the discovery of penicillin from the fungus *Penicillium* [9]. In recent years, a wealth of novel fungal biosynthetic pathways and compounds have been identified [10–15], suggesting fungi remain a viable source for developing new antibiotics.

Manaaki Whenua, a Crown Research Institute in Aotearoa New Zealand, is the custodian of the International Collection of Microorganisms from Plants (ICMP), which contains thousands of fungi and bacteria primarily sourced from Aotearoa and the South Pacific [16]. While the collection includes some fungal genera traditionally used for antibiotic production, it has not been rigorously tested for antimicrobial activity. We have previously reported the results of our efforts to screen a small portion of the collection for anti-mycobacterial activity [17]. Of relevance, we identified several ICMP fungal isolates with activity against *Mycobacterium abscessus* and *M. marinum*, including an unknown species of *Boeremia* and an isolate of a novel genus and species in the family Phanerochaetaceae [17].

Here, we use a ‘one strain-many compounds’ (OSMAC) approach [18, 19] – growing the fungi on seven different media – to screen 32 ICMP isolates for activity against *E. coli*. Our results indicate that several of the tested fungi possess anti-*E. coli* activity and are suitable for further study, including the aero-aquatic Ascomycete *Hyaloscypha* sp. ICMP 16864.

## Materials and Methods

### Bacterial strains and growth conditions

In this study, we used a bioluminescent derivative of the antibiotic-testing strain *E. coli* ATCC 25922, designated 25922 lux [20], expressing the bacterial luciferase (*lux*) operon from the integrating plasmid p16Slux [21]. We grew *E. coli* 25922 lux cultures in Mueller Hinton media (Fort Richard, New Zealand) at 37 °C, shaking at 200 rpm as necessary.

### Fungal material and growth conditions

Fungal isolates (Table 1) were provided by Manaaki Whenua—Landcare Research, the New Zealand Crown Research Institute responsible for curating the International Collection of Microorganisms from Plants (ICMP). As previously described, we stored fungal isolates individually in cryotubes at -80 °C [17]. Briefly, we made freezer stocks by growing each fungus on 1.5% Potato Dextrose Agar (PDA) at room temperature (approx. 20 °C) and excising small cubes of agar (5–6 mm in length) from the fungus’ growing edge. We placed these cubes within a cryovial containing 1 mL of 10% glycerol and rested them for one hour, after which we removed the remaining liquid glycerol and stored the tubes at −80 °C.

**Table 1.**
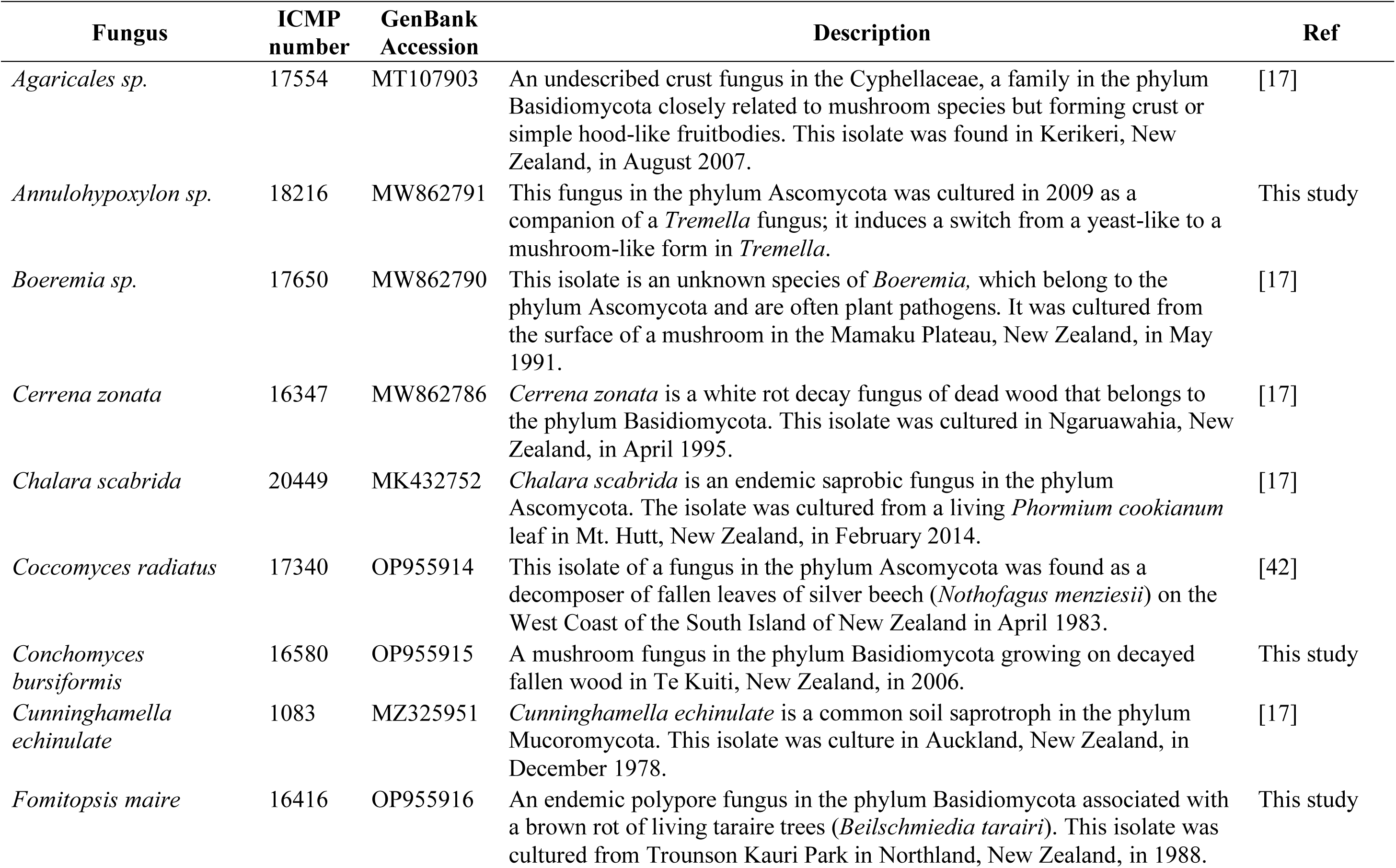
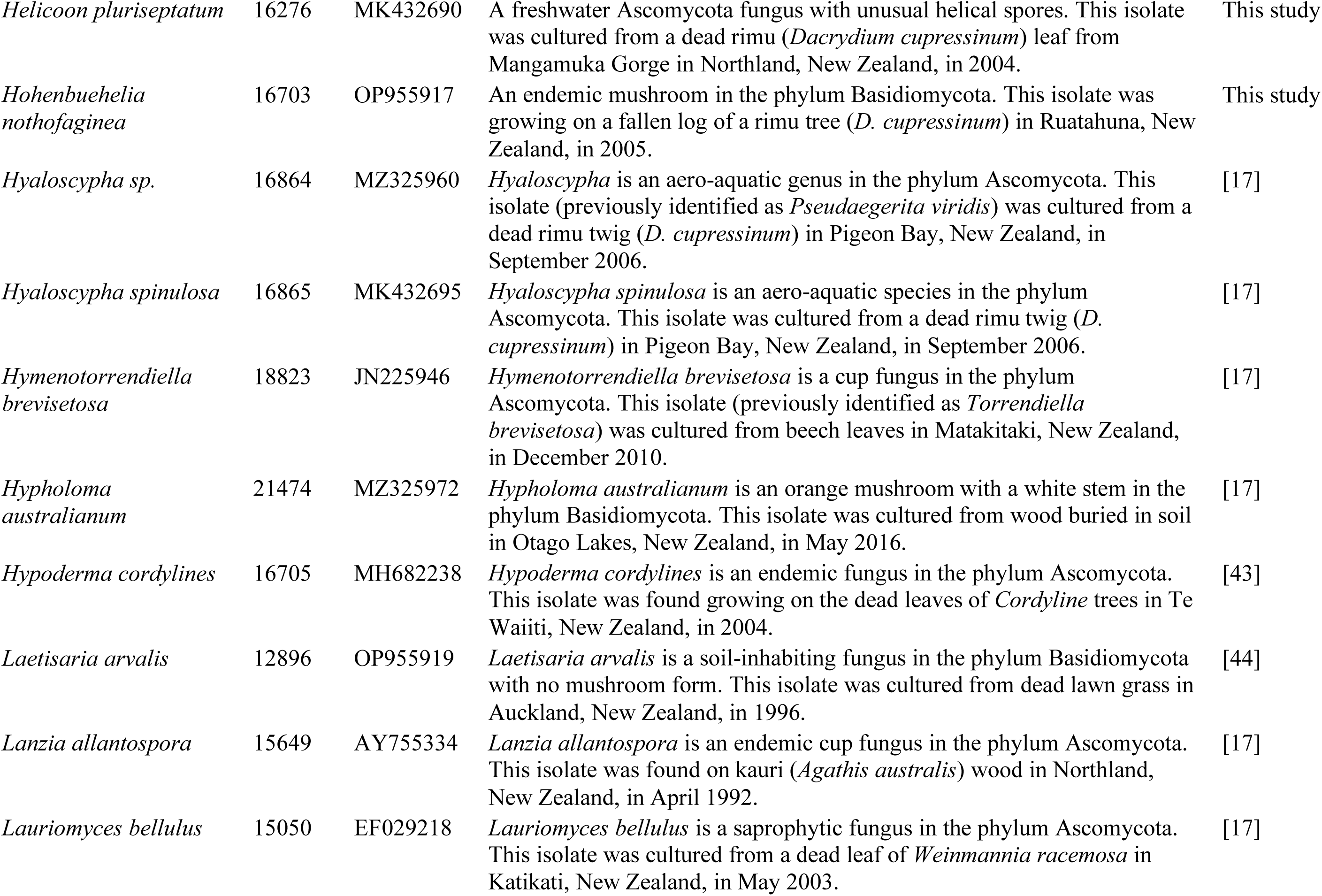
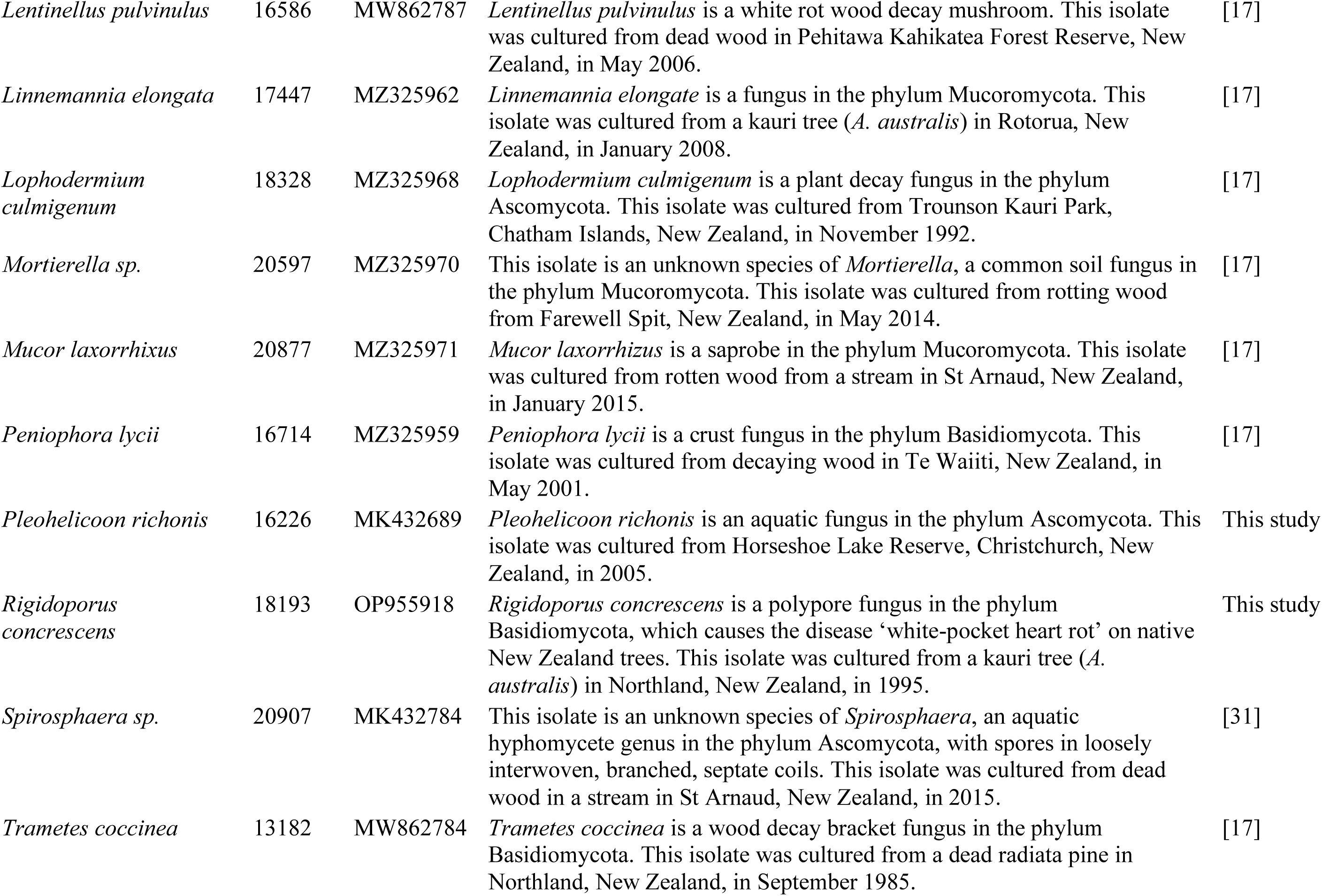
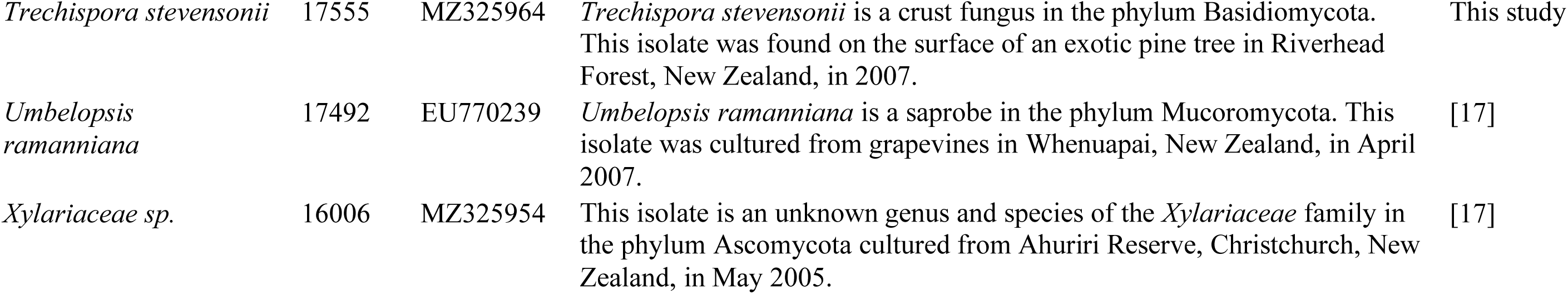
Fungal isolates used in this study.

### Fungal DNA extraction and ITS sequencing

As previously described [17], we used a small portion of mycelium from growing fungi and extracted DNA using the REDExtract-N-Amp^TM^ Plant PCR ReadyMix (Sigma-Aldrich, New Zealand) according to the manufacturer’s protocol. Briefly, we diluted DNA samples five-fold and amplified them using the ITS1F (5’ CTTGGTCATTTAGAGGAAGTAA 3’) and ITS4 (5’ TCCTCCGCTTATTGATATGC 3’) primer set in a 10 µl reaction. We used the following PCR conditions: initial denaturation at 94 °C for three minutes, followed by 40 cycles of denaturation at 94 °C for 30 s, annealing at 52 °C for 30 s and extending at 72 °C for 30 s. The final extension was performed at 72 °C for seven minutes. We checked the amplified DNA by gel electrophoresis before sequencing using an Applied Biosystems™ 3500xL Genetic Analyzer using ITS1F and ITS4 primers. We trimmed and combined the sequence data using Geneious Prime (Biomatters, New Zealand), removed any low-quality reads and used BLAST to check fungal identification. Optimised sequence data were aligned using MEGA7 [22].

### Primary fungal screening using the zone of inhibition assay

We first grew each fungal isolate on Potato Dextrose Agar (PDA). Once grown, we excised 1cm cubes from the growing edge using a scalpel and placed single cubes onto the centre of 90 mm Petri dishes containing either PDA or one of six other growth media: Czapek Solution Agar (CSA), Czapek Yeast Extract Agar (CYA), Malt Extract Agar (MEA), Oatmeal Agar (OA), Rice Extract Agar (REA), and Water Agar (WA). We measured the diameter of each fungus at regular intervals over 60 days (Fig. 1). All fungal isolates were incubated at room temperature.

**Figure 1.**
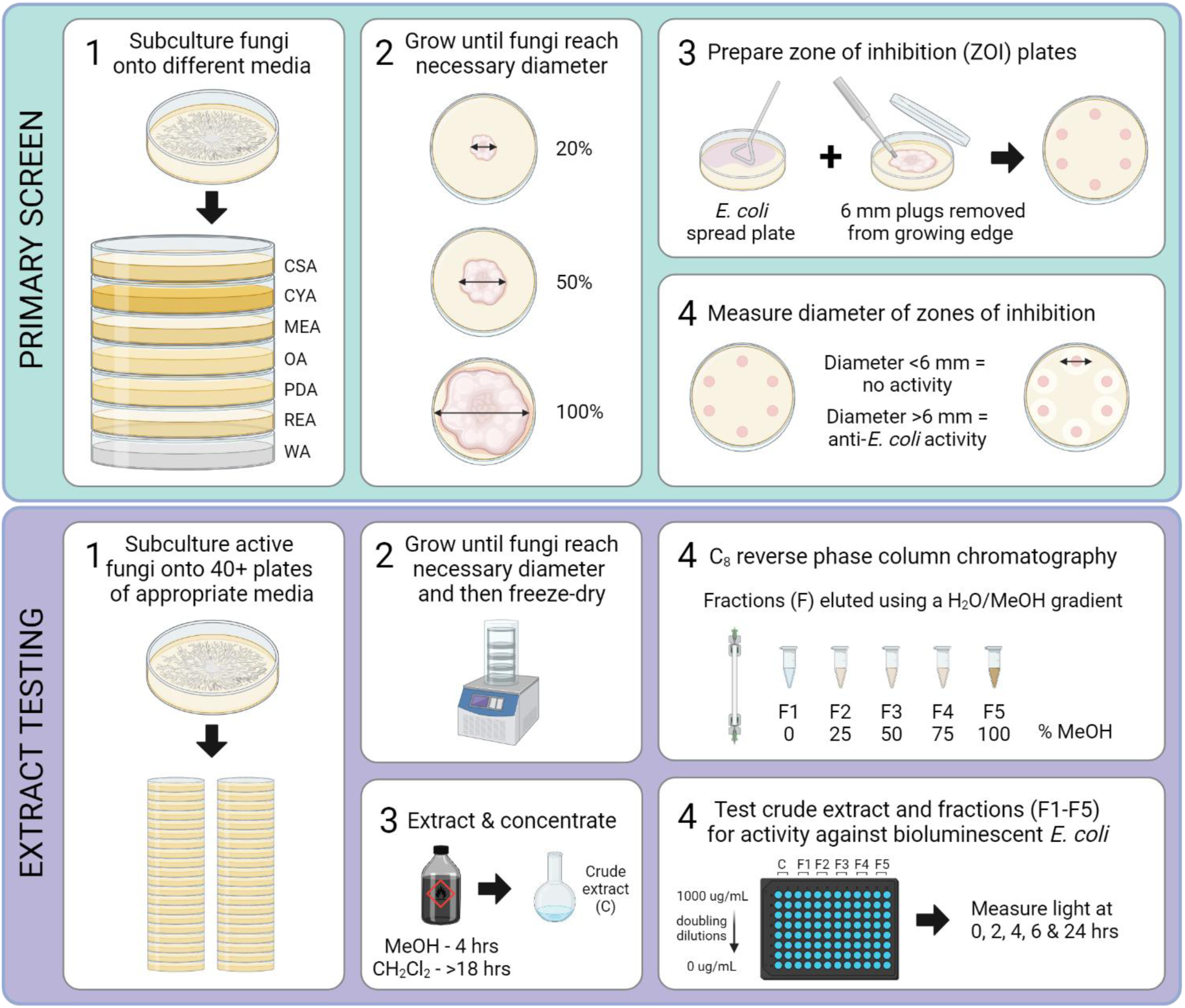
Schematic of experimental workflow. Schematic produced using Biorender.com.

Where possible, we tested each fungus for anti-*E. coli* activity once they had grown to cover 20, 50, and 100 % of a 90 mm Petri dish. We used a modified version of the zone of inhibition (ZOI) assay, which we have published on the protocol repository website protocols.io [23]. Briefly, overnight cultures of *E. coli* 25922 lux were diluted in MHB to an optical density at 600 nm (OD_600_) of 0.01 (the equivalent of ∼10^6^ bacteria per mL) and used to inoculate CSA, CYA, MEA, OA, PDA, REA, and WA plates with a lawn of *E. coli*. Using a 6 mm punch biopsy tool (Paramount, Capes Medical, New Zealand), we removed plugs from the growing edge of each fungus and placed these fungus side down onto the corresponding *E. coli* lawn plates. We incubated these plates overnight at 37 °C, after which we measured the diameter of any zones of inhibition (Fig. 1). We performed these assays using at least three independent biological replicates of each fungus.

### General chemistry conditions

We recorded NMR spectra using a Bruker Avance DRX-400 spectrometer or an Avance III-HD 500 spectrometer operating at 400 MHz or 500 MHz for ^1^H nuclei and 100 MHz or 125 MHz for ^13^C nuclei utilising standard pulse sequences at 298 K. We recorded high resolution mass spectra on a Bruker micrOTOF QII (Bruker Daltonics, Bremen, Germany). We carried out analytical thin layer chromatography (TLC) on 0.2 mm thick plates of DC-plastikfolien Kieselgel 60 F254 (Merck, New Zealand). We carried out reversed-phase column chromatography on C_8_ support with a pore size of 40-63 µm (Merck, New Zealand). We performed gel filtration chromatography on Sephadex LH-20 (Pharmacia, New Zealand). We carried out flash chromatography on Diol-bonded silica with a pore size of 40–63 microns (Merck, New Zealand). We used solvents of analytical grade or better and/or purified according to standard procedures.

### Fungal fermentation and extraction

We grew fungal cultures on solid media at room temperature and then freeze-dried them (Fig. 1). We extracted the dry cultures with MeOH (Sigma-Aldrich, New Zealand) for four hours, followed by CH_2_Cl_2_ (Sigma-Aldrich, New Zealand) overnight. We concentrated the combined organic extracts under reduced pressure. We subjected the crude extracts to C8 reversed-phase column chromatography eluting with a gradient of H_2_O/MeOH (Sigma-Aldrich, New Zealand) to afford five fractions (F1–F5) (Fig. 1). We have published our protocol on the website protocols.io (van de Pas and Wiles, 2023c). Full details are provided in Supplementary Methods. 53 MEA plates were inoculated with ICMP 16347 and incubated at room temperature for ten days. Fully grown fungal plates were freeze-dried (67.33 g, dry weight) and extracted with MeOH (1 L) for four hours, followed by CH_2_Cl_2_ (1 L) overnight. Combined organic extracts were concentrated under reduced pressure to afford brown oil (4.76 g). The crude product was subjected to C_8_ reversed-phase column chromatography eluting with a gradient of H_2_O/MeOH to afford five fractions (F1–F5). F4 was purified by diol-bonded silica gel column chromatography, eluting with gradient n-hexane/EtOAc to afford four fractions (A1–A4). Further purification of fraction A1 by silica gel column chromatography, eluting with gradient n-hexane/EtOAc, afforded steperoxide A (**2**) (30.13 mg). Purification of A2 by silica gel column chromatography, eluting with gradient n-hexane/EtOAc, afforded merulin A (**1**) (7.16 mg).

19 CYA plates were inoculated with ICMP 16347 and incubated at room temperature for ten days. Fully grown fungal plates were freeze-dried (15.88 g, dry weight) and extracted with MeOH (400 mL) for four hours, followed by CH_2_Cl_2_ (400 mL) overnight. Combined organic extracts were concentrated under reduced pressure to afford a brown oil (2.70 g). The crude product was subjected to C_8_ reversed-phase column chromatography eluting with a gradient of H_2_O/MeOH to afford five fractions (F1–F5). F3 was subjected to purification by Sephadex LH-20, eluting with MeOH, to afford seven fractions (A1–A7). Further purification of fraction A3 by silica gel column chromatography, eluting with gradient n-hexane/EtOAc, afforded antroalbol H (**3**) (4.62 mg). Purification of A2 by silica gel column chromatography, eluting with gradient n-hexane/EtOAc, followed by trituration with methanol afforded merulin A (**1**) (9.90 mg) and steperoxide A (**2**) (3.54 mg).

37 CYA plates were inoculated with ICMP 17650 and incubated at room temperature for ten days. Fully grown fungal plates were freeze-dried (28.68 g, dry weight) and extracted with MeOH (750 mL) for four hours, followed by CH_2_Cl^2^ (750 mL) overnight. Combined organic extracts were concentrated under reduced pressure to afford an orange oil (4.00 g). The crude product was subjected to C_8_ reversed-phase column chromatography eluting with a gradient of H_2_O/MeOH to afford five fractions (F1–F5). F4 was triturated with dichloromethane, and the precipitate was purified by silica gel column chromatography, eluting with gradient n-hexane/EtOAc, to afford cytochalasin B (**4**) (23.19 mg).

### Minimum inhibitory concentration testing of fungal extracts

We have published a detailed description of our method on the protocol repository website protocols.io [24]. Briefly, we grew *E. coli* 25922 lux overnight. Then, we diluted the culture in Mueller Hinton broth II (MHB) (Fort Richard, New Zealand) to give an optical density at 600 nm (OD_600_) of 0.001, which is the equivalent of ∼5 x 10^5^ bacteria per mL. We dissolved the fungal fractions in DMSO (Sigma-Aldrich, New Zealand). We added these in duplicate to the wells of a black 96-well plate (Nunc, Thermo Scientific) at doubling dilutions with a maximum concentration of 2000 μg/mL (Fig. 1). We then added 50 μL of diluted *E. coli* 25922 lux to each well giving final extract concentrations of 0-1000 μg/mL and a cell density of ∼10^5^ CFU/mL. We used the antibiotic erythromycin (Sigma-Aldrich, New Zealand) as a positive control at 250 μg/mL. We measured bacterial luminescence at 0, 2, 4, 6, and 24 hours using a Victor X-3 luminescence plate reader (PerkinElmer) with an integration time of 1 s. Between measurements, plates were covered, placed in a plastic box lined with damp paper towels, and incubated with shaking at 100 rpm at 37 °C. We have defined the MIC as causing a 1 log reduction in light production, as previously described [25, 26].

### Antimicrobial testing of pure compounds

Antimicrobial evaluation of the pure compounds against *Acinetobacter baumannii* ATCC 19606, *Candida albicans* ATCC 90028, *Cryptococcus neoformans* ATCC 208821, *E. coli* ATCC 25922, *Klebsiella pneumoniae* ATCC 700603, *Pseudomonas aeruginosa* ATCC 27853, and methicillin-resistant *Staphylococcus aureus* ATCC 43300 (MRSA) was undertaken at the Community for Open Antimicrobial Drug Discovery at The University of Queensland (Queensland, Australia) according to standard protocols [27]. As previously described [13], bacterial strains were cultured in either Luria broth (LB) (In Vitro Technologies, USB75852, Victoria, Australia), nutrient broth (NB) (Becton Dickson, 234,000, New South Wales, Australia) or MHB at 37 °C overnight [27]. A sample of culture was then diluted 40-fold in fresh MHB and incubated at 37 °C for 1.5−2 h. The compounds were serially diluted 2-fold across the wells of 96-well plates (Corning 3641, nonbinding surface), with compound concentrations ranging from 0.015 to 64 μg/mL, plated in duplicate. The resultant mid-log phase cultures were diluted to the final concentration of 1 × 10^6^ CFU/mL. Then, 50 μL was added to each well of the compound-containing plates, giving a final compound concentration range of 0.008– 32 μg/mL and a cell density of 5 × 10^5^ CFU/mL. All plates were then covered and incubated at 37 °C for 18 hours. Resazurin was added at 0.001% final concentration to each well and incubated for two hours before MICs were read by eye.

*Candida albicans* and *Cryptococcus neoformans* were cultured for three days on Yeast Peptone Dextrose (YPD) agar (Becton Dickinson, 233,520, New South Wales, Australia) at 30 °C. A yeast suspension of 1 × 10^6^ to 5 × 10^6^ CFU/mL was prepared from five colonies. These stock suspensions were diluted with Yeast Nitrogen Base (YNB) (Becton Dickinson, 233,520, New South Wales, Australia) broth to a final concentration of 2.5 × 10^3^ CFU/mL. The compounds were serially diluted 2-fold across the wells of 96-well plates (Corning 3641, nonbinding surface), with compound concentrations ranging from 0.015 to 64 μg/mL and final volumes of 50 μL, plated in duplicate. Then, 50 μL of a previously prepared fungi suspension, in YNB broth to the final concentration of 2.5 × 10^3^ CFU/mL, was added to each well of the compound-containing plates, giving a final compound concentration range of 0.008–32 μg/mL. Plates were covered and incubated at 35 °C for 36 h without shaking. *Candida albicans* MICs were determined by measuring optical density at 530 nm. For *Cryptococcus neoformans*, resazurin was added at 0.006% final concentration to each well and incubated for three hours before MICs were determined by measuring the optical density at 570–600 nm.

Colistin and vancomycin were used as positive bacterial inhibitor standards for Gram-negative and Gram-positive bacteria, respectively. Fluconazole was used as a positive fungal inhibitor standard for *Candida albicans* and *Cryptococcus neoformans*. The antibiotics were provided in four concentrations, with two above and two below their MIC value, and plated into the first eight wells of Column 23 of the 384-well NBS plates. The quality control of the assays was determined by the antimicrobial controls and the Z’-factor (using positive and negative controls). Each plate was deemed to fulfil the quality criteria if the Z’-factor was above 0.4, and the antimicrobial standards showed a full range of activity, with total growth inhibition at their highest concentration and no growth inhibition at their lowest concentration [27].

### Statistical analysis

After confirming that residuals were approximately normally distributed and checking that a random intercept was sufficient, we fitted mixed ANOVA models for log (size) with a random intercept for biologic replicates within organisms. Two observations of zero size were omitted. We tested the main effects and second-order interactions of the variables using the lme4 and car packages in R [28–30].

## Results

### Identification of a novel fungus endemic to Aotearoa New Zealand

The 32 ICMP fungal isolates used in this study belong to three different phyla: the Ascomycota (n=15), the Basidiomycota (n=12), and the Mucoromycota (n=5) (Table 1) and were collected between 1978 and 2016 (Table 1) from locations across Aotearoa New Zealand, including the North, South, and Chatham Islands (Fig. 2). Of the 32 isolates, six are of unknown species (Table 1) and we and others have previously reported five of them [17, 31]. The remaining unknown species is represented by ICMP 18216, which belongs to the *Annulohypoxylon* genus and was isolated in 2009 as a companion of a *Tremella* fungus (Table 1).

**Figure 2.**
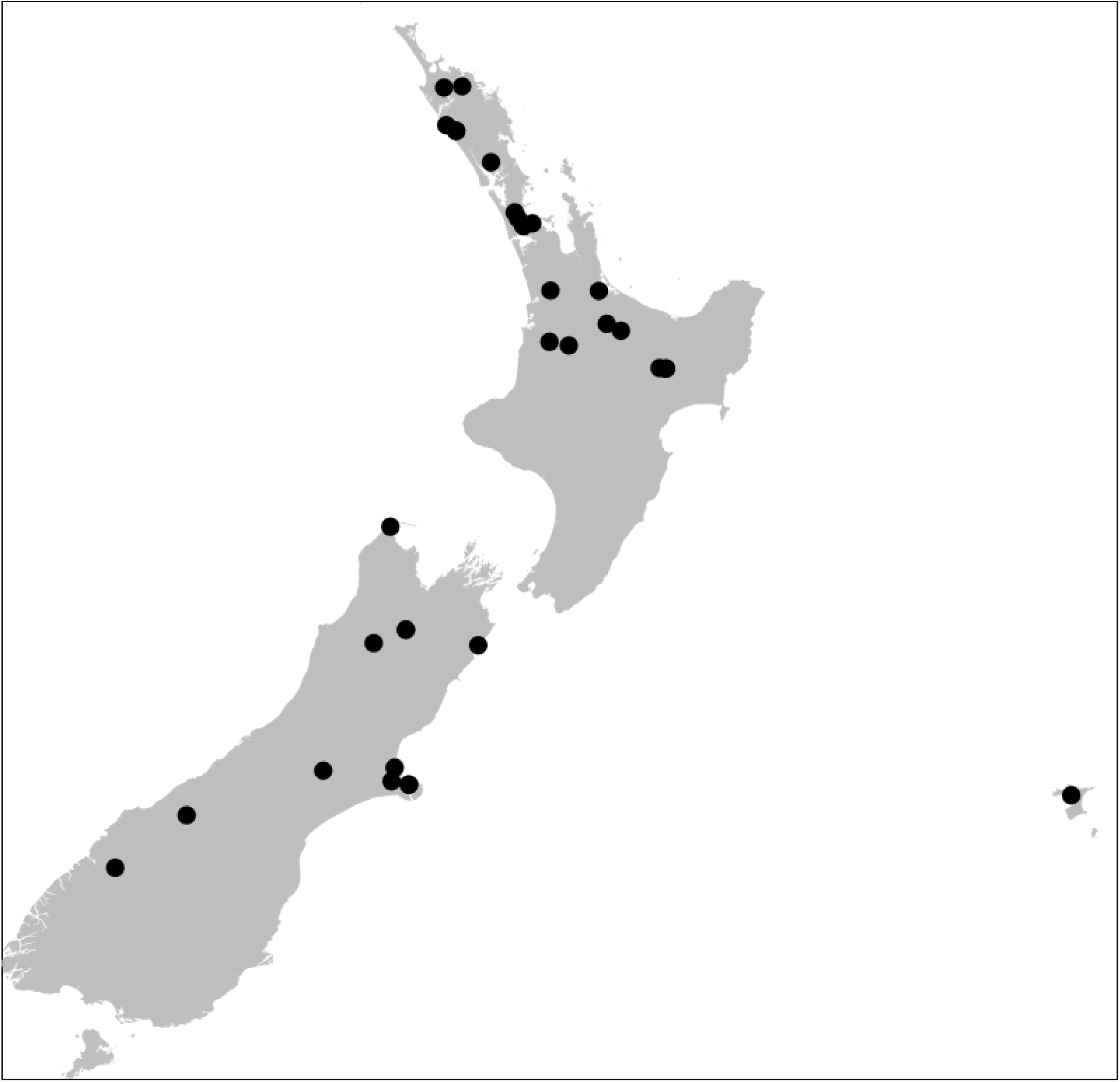
Geographical spread of isolation locations within the archipelago of Aotearoa New Zealand for the fungi used in this study. Black dots are individual ICMP fungal isolates.

### Growth characteristics of a panel of ICMP fungal isolates

We assessed the growth of the 32 ICMP isolates on seven different media: Czapek Solution Agar (CSA), Czapek Yeast Extract Agar (CYA), Malt Extract Agar (MEA), Oatmeal Agar (OA), Potato Dextrose Agar (PDA), Rice Extract Agar (REA), and Water Agar (WA), measuring the diameter of each fungus at regular intervals over 60 days (Fig. 3).

**Figure 3.**
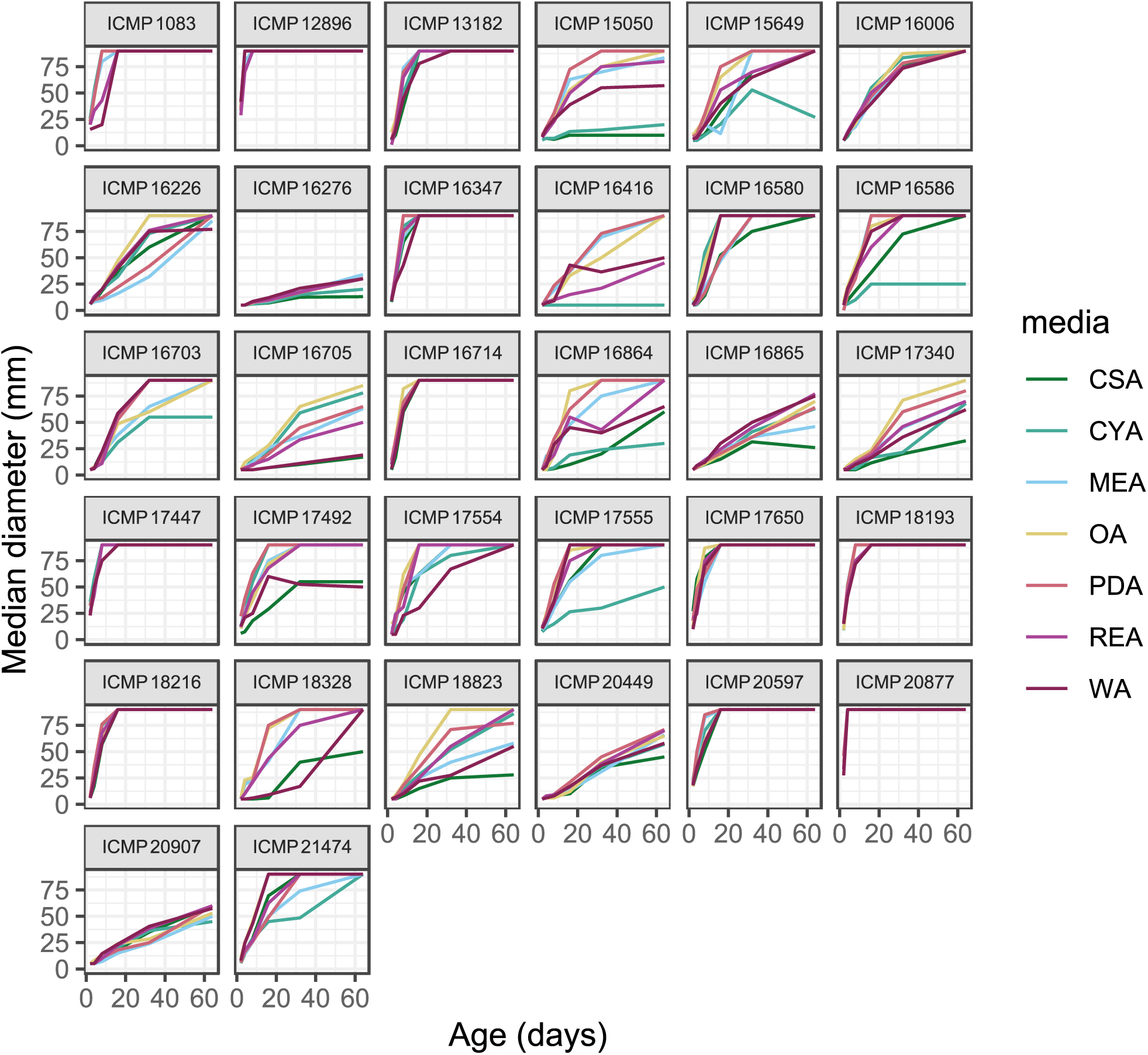
Growth of ICMP isolates over 60 days on seven different media. Data are presented as median size in mm of at least three biological replicates. Media: CSA, Czapek Solution Agar; CYA, Czapek Yeast extract Agar; MEA, Malt Extract Agar; OA, Oatmeal Agar; PDA, Potato Dextrose Agar; REA, Rice Extract Agar; WA, Water Agar.

We observed that the ICMP isolates could be divided into four categories: 1) those that grew to entirely cover a 90 mm Petri dish by 16 days regardless of the growth medium (10/32 [31.3%]) (Fig. 4); 2) those that grew to entirely cover a 90 mm Petri dish by 60 days regardless of the growth medium (4/32 [12.5%]) (Fig. 3); 3) those that grew variably, entirely covering a 90 mm Petri dish by 60 days when grown on at least one medium (13/32 [40.6%]) (Fig. 3); and 4) those that did not entirely cover a 90 mm Petri dish by 60 days when grown on any of the seven media (5/32 [15.6%]) (Fig. 3). The Mucoromycota were over-represented in the first category with 4/5 of the isolates able to grow rapidly on all the media tested, while the Ascomycota were over-represented in the fourth category (5/5) comprising those ICMP isolates that did not grow to cover a 90 mm Petri dish within 60 days on any media.

**Figure 4.**
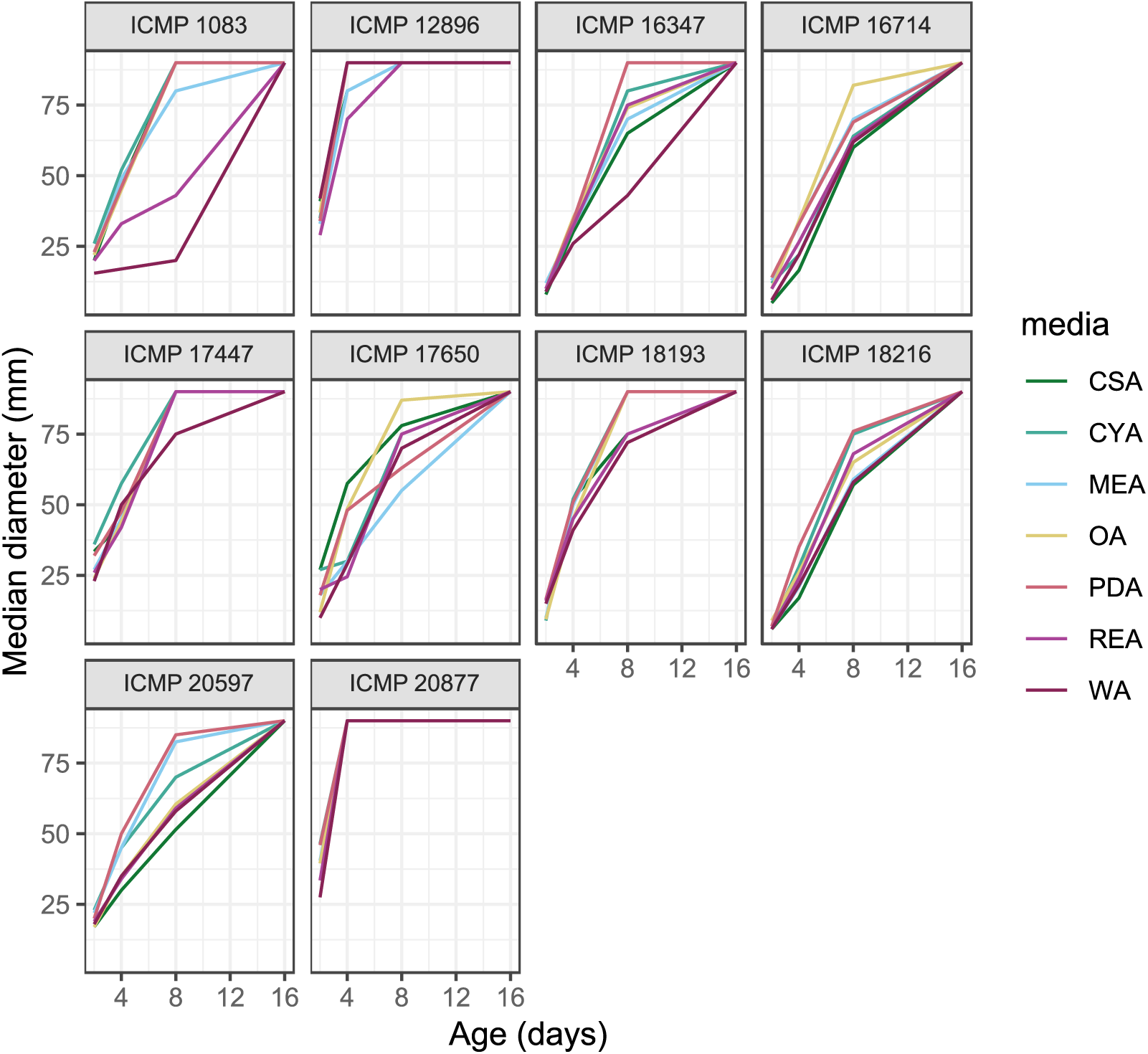
Growth of fast-growing ICMP isolates on seven different media. Data are presented as median size in mm of at least three biological replicates. Media: CSA, Czapek Solution Agar; CYA, Czapek Yeast extract Agar; MEA, Malt Extract Agar; OA, Oatmeal Agar; PDA, Potato Dextrose Agar; REA, Rice Extract Agar; WA, Water Agar.

We calculated the growth rate (as mm/day) of each ICMP isolate when actively growing on each of the seven media (Table 2, Figs. 3, 4). We observed strong statistical evidence of differences in growth rate between phyla (p=0.000125) and between media (p=<2.2x10^-16^), and of interactions between phylum and medium (p=< 2.671x10^-16^). Some isolates grew as slowly as 1 mm/day on some media (for example, *Fomitopsis maire* ICMP 16416, *Helicoon pluriseptatum* ICMP 16276, *Hypoderma cordylines* ICMP 16705, and *Lophodermium culmigenum* ICMP 18328), while others grew at a rate of more than 20 mm/day (*Laetisaria arvalis* ICMP 12896 and *Mucor laxorrhixus* ICMP 20877) (Table 2). On average, the Mucoromycota isolates grew faster than the Basidiomycota isolates, which were faster than the Ascomycota. Median growth rates for the Mucoromycota isolates ranged from 8.1 to 12.9 mm/day depending on the growth medium, while for the Basidiomycota and Ascomycota, they were 3.8 to 5.9 and 1.2 to 1.4 mm/day, respectively (Table 2).

**Table 2.** Fungal growth rates (mm/day) on various solid media.

| Media<sup>1</sup> | All ICMP isolates (n=32) |  |  | Ascomycota (n=15) |  |  | Basidiomycota (n=12) |  |  | Mucoromycota (n=5) |  |  |
| --- | --- | --- | --- | --- | --- | --- | --- | --- | --- | --- | --- | --- |
|  | Min. | Max. | Median | Min. | Max. | Median | Min. | Max. | Median | Min. | Max. | Median |
| CSA | 0.7<br>(ICMP<br>16705) | 19.5<br>(ICMP<br>12896) | 2.4 | 0.7<br>(ICMP<br>16705) | 11.8<br>(ICMP<br>17650) | 1.2 | 0.9<br>(ICMP<br>16416) | 19.5<br>(ICMP<br>12896) | 3.8 | 2.2<br>(ICMP<br>17492) | 19.5<br>(ICMP<br>20877) | 10.8 |
| CYA | 0.8<br>(ICMP<br>16276) | 23.3<br>(ICMP<br>20877) | 2.7 | 0.8<br>(ICMP<br>16276) | 9.6<br>(ICMP<br>17650) | 1.2 | 0.9<br>(ICMP<br>16416) | 19.4<br>(ICMP<br>12896) | 3.8 | 6.9<br>(ICMP<br>17492) | 23.3<br>(ICMP<br>20877) | 12.9 |
| MEA | 0.8<br>(ICMP<br>16276) | 19.8<br>(ICMP<br>20877) | 3.5 | 0.8<br>(ICMP<br>16276) | 7.9<br>(ICMP<br>17650) | 1.3 | 2.5<br>(ICMP<br>16580) | 18.2<br>(ICMP<br>12896) | 4.2 | 6.0<br>(ICMP<br>17492) | 19.8<br>(ICMP<br>20877) | 10.9 |
| OA | 0.9<br>(ICMP<br>16276) | 19.6<br>(ICMP<br>12896) | 4.7 | 0.9<br>(ICMP<br>16276) | 9.2<br>(ICMP<br>17650) | 1.6 | 1.6<br>(ICMP<br>16416) | 19.6<br>(ICMP<br>12896) | 5.9 | 5.1<br>(ICMP<br>17492) | 18.6<br>(ICMP<br>20877) | 11.2 |
| PDA | 0.8<br>(ICMP<br>16276) | 22.8<br>(ICMP<br>20877) | 4.0 | 0.8<br>(ICMP<br>16276) | 10.7<br>(ICMP<br>17650) | 1.5 | 2.4<br>(ICMP<br>16580) | 19.8<br>(ICMP<br>12896) | 5.5 | 9.4<br>(ICMP<br>17492) | 22.8<br>(ICMP<br>20877) | 11.6 |
| REA | 0.8<br>(ICMP<br>16276) | 16.4<br>(ICMP<br>20877) | 3.5 | 0.8<br>(ICMP<br>16276) | 9.1<br>(ICMP<br>17650) | 1.4 | 1.4<br>(ICMP<br>16416) | 15.2<br>(ICMP<br>12896) | 4.8 | 5.6<br>(ICMP<br>17492) | 16.4<br>(ICMP<br>20877) | 8.3 |
| WA | 0.7<br>(ICMP<br>18328) | 20.5<br>(ICMP<br>12896) | 3.7 | 0.7<br>(ICMP<br>18328) | 7.6<br>(ICMP<br>17650) | 1.2 | 1.7<br>(ICMP<br>16416) | 20.5<br>(ICMP<br>12896) | 4.9 | 4.7<br>(ICMP<br>17492) | 13.9<br>(ICMP<br>20877) | 8.1 |

Key: <sup>1</sup>Czapek Solution Agar (CSA); Czapek Yeast Extract Agar (CYA); Malt Extract Agar (MEA); Oatmeal Agar (OA); Potato Dextrose Agar (PDA); Rice Extract Agar (REA); Water Agar (WA).

### Whole-cell screening identified many ICMP fungal isolates as having anti-*E. coli* activity

We screened 32 ICMP fungal isolates grown on seven different media for antibacterial activity against *E. coli* ATCC 25922 lux. Where possible, we tested each fungus for activity once it had grown to cover 20, 50, and 100 % of a 90 mm Petri dish. We measured antibacterial activity as the production of zones of inhibition after incubation with *E. coli* ATCC 25922 lux for 24 hours. As the fungal plugs were 6 mm in diameter, a diameter greater than 6 mm indicates the production of a zone of inhibition and, therefore, anti-*E. coli* activity (Fig. 5). We observed strong statistical evidence of the presence of a zone of inhibition differing by phylum (p=0.0032) and media (p=0.0014) and by the combination of the two (p=0.041). There was no evidence of a difference by fungal age.

**Figure 5.**
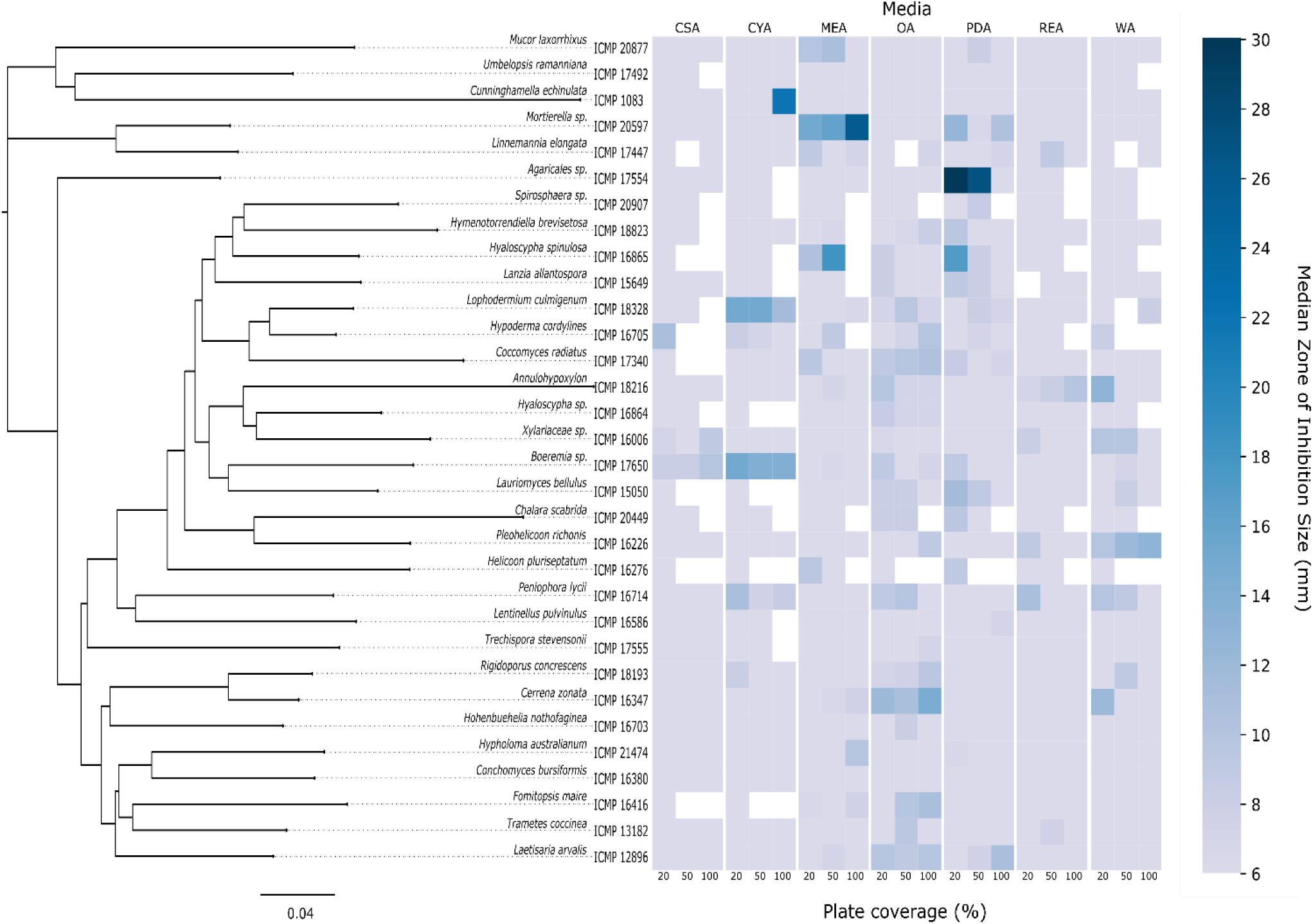
Phylogeny of ICMP isolates and their production of zones of inhibition against *E. coli* when grown on different media. Key: Isolates were tested for activity against *E. coli* ATCC 25922 lux in zone of inhibition assays when they had grown to cover 20, 50, and 100% of the area of a Petri dish. Data are presented as median zones of inhibition (mm radius) of at least three biological replicates. Media: CSA, Czapek Solution Agar; CYA, Czapek Yeast extract Agar; MEA, Malt Extract Agar; OA, Oatmeal Agar; PDA, Potato Dextrose Agar; REA, Rice Extract Agar; WA, Water Agar. The phylogenetic tree was constructed in GeneiousPrime (v2020.0.5) (Biomatter Inc.) using the neighbour-joining and Tamura-Mei genetic distance models. The tree was constructed by comparing ITS sequences to those in GenBank, and consensus sequences assembled de-novo using Geneious assembler.

### Anti-E. coli activity varies by fungal phylum

We converted the diameter of the zone of inhibition (ZOI) produced by the fungal isolates into ZOI scores. A diameter of 6 mm, the size of the fungal plug, is given a ZOI score of 0. Zones with a diameter of 7-10 mm are scored as 1, 11-15 mm diameters as 2, and zones with a diameter of >15 mm are scored as 3.

If we define a fungus-medium combination as being active against *E. coli* if the median ZOI score is 1 or higher, then only two fungal isolates, ICMP 17492 (*Umbelopsis ramanniana*) and ICMP 20907 (*Spirosphaera sp.*), were not active (Fig. 6). The phylum with the most active fungus-medium combinations was the Ascomycota (46/105 [43.8%]), followed by the Basidiomycota (28/84 [33.3%]) and then the Mucoromycota (7/35 [20.0%]) (Fig. 6).

**Figure 6.**
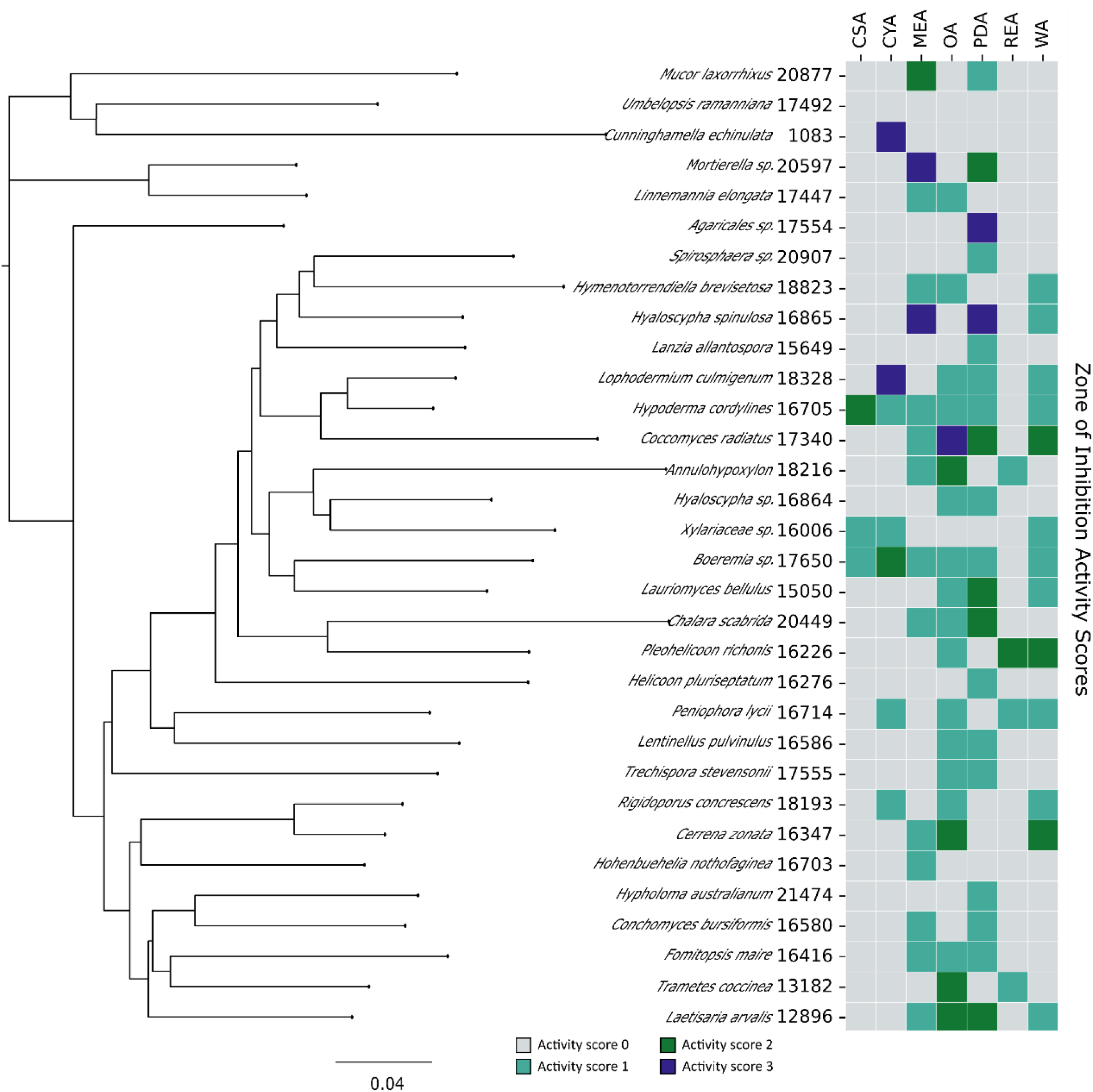
Phylogeny of ICMP isolates and their anti-*E. coli* activity when grown on different media. Isolates were tested for activity against *E. coli* ATCC 25922 lux, and zones of inhibition (ZOI) data were converted into ZOI scores. A score of 0 represents no activity, a score of 1 equates to the production of zones with a diameter of 7-10 mm, a score of 2 with zones with diameters of 11-15 mm, and a score of 3 to zones with a diameter of >15 mm. Data are presented as median activity scores of at least three biological replicates. Media: CSA, Czapek Solution Agar; CYA, Czapek Yeast extract Agar; MEA, Malt Extract Agar; OA, Oatmeal Agar; PDA, Potato Dextrose Agar; REA, Rice Extract Agar; WA, Water Agar. The phylogenetic tree was constructed in GeneiousPrime (v2020.0.5) (Biomatter Inc.) using the neighbour-joining and Tamura-Mei genetic distance models. The tree was constructed by comparing ITS sequences to those in GenBank, and consensus sequences assembled de-novo using Geneious assembler.

If we use a more stringent measure of activity, with a fungal isolate required to produce a zone of inhibition with a diameter of greater than 10 mm (a ZOI score ≥2), then the number of active ICMP isolates reduces from 30 to 16 (Fig. 6). The Ascomycota and Mucoromycota have a similar percentage of active fungus-medium combinations, 12.4% (13/105) and 11.4% (4/35), respectively, while the Basidiomycota are the group with the least percentage of active combinations (7.1% [6/84]).

Six ICMP isolates produced zones of inhibition with a diameter of >15 mm (corresponding to a ZOI score of 3): the Ascomycota *Coccomyces radiatus* (ICMP 17340), *Hyaloscypha spinulosa* (ICMP 16865), and *Lophodermium culmigenum* (ICMP 18328); the Basidiomycete *Agaricales sp.* (ICMP 17554); and the Mucoromycota *Cunninghamella echinulate* (ICMP 1083) and *Mortierella sp.* (ICMP 20597) (Fig. 6).

### Differential impact of growth medium on anti-E. coli activity

We observed that many of the ICMP fungi displayed differential activity depending on their growth medium (Figs. 5, 6). Using a ZOI score ≥1, 24/30 ICMP isolates were active on more than one medium, with seven isolates displaying activity when grown on four or more media. Using A ZOI score ≥2, 6/16 ICMP isolates were active on two or more media. The growth medium which resulted in the least anti-*E. coli* activity was Czapek Solution Agar (CSA). In contrast, growth on Oatmeal Agar (OA) and Potato Dextrose Agar (PDA) resulted in the most activity, with 19/30 and 20/30 ICMP isolates, respectively, exhibiting a ZOI score ≥1. This was true when using the more stringent ZOI score of ≥2 (Fig. 6).

### Screening of extracts and fractions from ICMP fungal isolates for anti-*E. coli* activity

Focusing on ICMP isolates which produced activity scores of 2 or above, we prepared extracts from 16 isolates comprising 20 fungus-medium combinations. We further separated these extracts into five fractions, designated F1-F5. As previously described, fraction F1 (100% water) is generally comprised of sugars, while fraction F5 (100% methanol) contains predominantly fatty acids and sterols [17]. Fractions F2, F3, and F4 typically contain the chemical compounds we are most interested in pursuing, with the potential to be bioactive.

We tested the crude extracts and fractions for activity against *E. coli* 25922 lux at concentrations ranging from 0-1000 µg/mL (Table 3, Fig. 7). We measured antibacterial activity as reductions in the light output of our bioluminescent *E. coli* strain over 24 hours and calculated activity scores as the negative log of the ratio of the AUC values of the fungus-containing measurements and the control measurements. An activity score above 1 corresponds to a >90% reduction in light compared to the control, while an activity score above 2 corresponds to a >99% reduction. We define an extract/fraction as active/antibacterial if the median activity score is above 1.

**Table 3.**
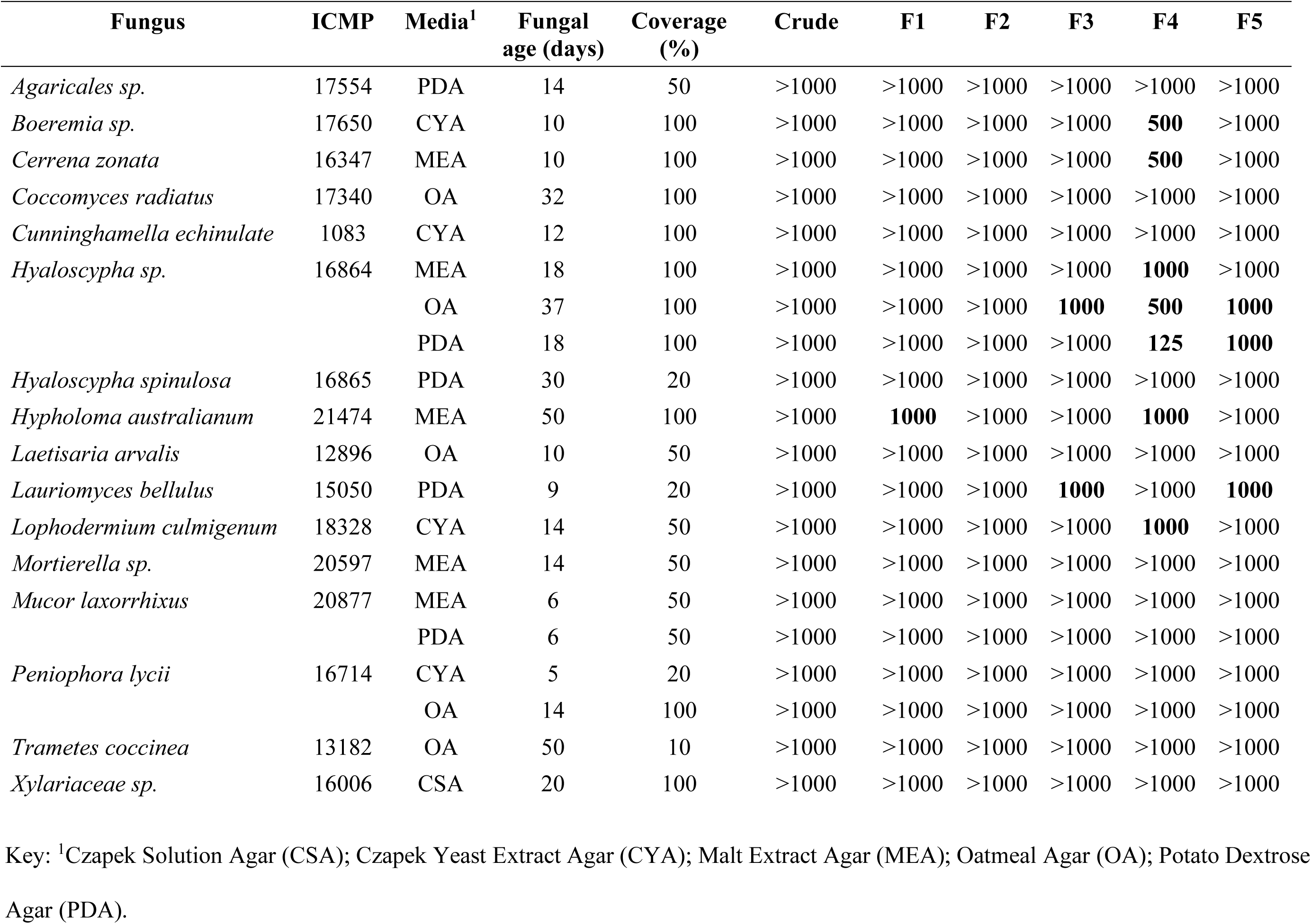
Minimum inhibitory concentrations (MIC) as µg/ml of ICMP fungal crude extracts and fractions (F1-5) against *E. coli*.

**Figure 7.**
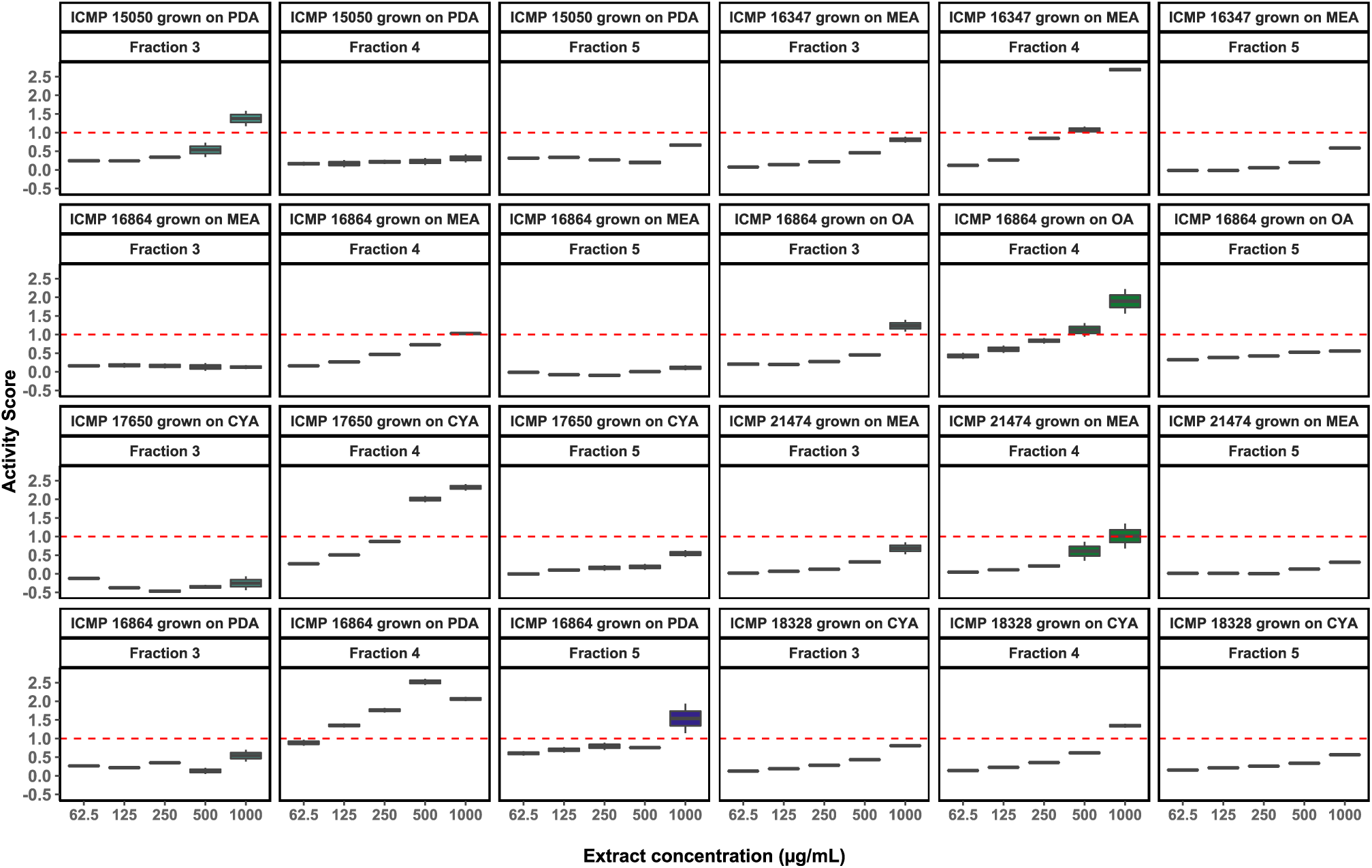
Antibacterial activity of fractions 3–5 from ICMP fungal isolates against *E. coli* 25922 lux. Data is presented as box and whisker plots of activity scores. The dotted line at 1 is the activity threshold. Scores above 1 correspond to a >90% reduction in bacterial bioluminescence compared to the corresponding no-fraction control. Similarly, an activity score above 2 corresponds to a >99% reduction. CYA, Czapek Yeast Extract Agar; MEA, Malt Extract Agar; OA, Oatmeal Agar; PDA, Potato Dextrose Agar. Boxes are upper and lower quartiles with the median shown. The whiskers extend up to 1.5× the inter-quartile range.

### Several ICMP fungal extracts and fractions retain anti-E. coli activity

Of the 20 fungus-medium combinations extracted and fractionated, extracts from 6 fungi retained activity with an MIC of at least 1000 µg/ml (Table 3). Of these, 3 had activity below 1000 µg/ml, all from fraction 4: *Boeremia* sp ICMP 17650 when grown on CYA (MIC of 500 µg/ml), *Cerrena zonata* ICMP 16347 when grown on MEA (MIC of 500 µg/ml), and *Hyaloscypha* sp. ICMP 16864 when grown on OA (MIC of 125 µg/ml) and PDA (MIC of 500 µg/ml) (Table 3, Fig. 7).

### Identification and activity testing of natural products from the active fractions obtained from *Boeremia sp*. ICMP 17650 and *Cerrena zonata* ICMP 16347

*Boeremia sp.* ICMP 17650 and *Cerrena zonata* ICMP 16347 were grown on CYA and MEA and extracted with methanol and di-chloromethane to afford crude extracts. The crude extracts were subjected to extensive chromatographic methods for purification, including C_8_ reversed-phase column chromatography (H_2_O/MeOH), Sephadex LH-20 (MeOH/5% CH_2_Cl_2_) and silica gel (hexane/EtOAc) chromatography to afford compounds **1**–**4** (Fig. 8).

**Figure 8.**
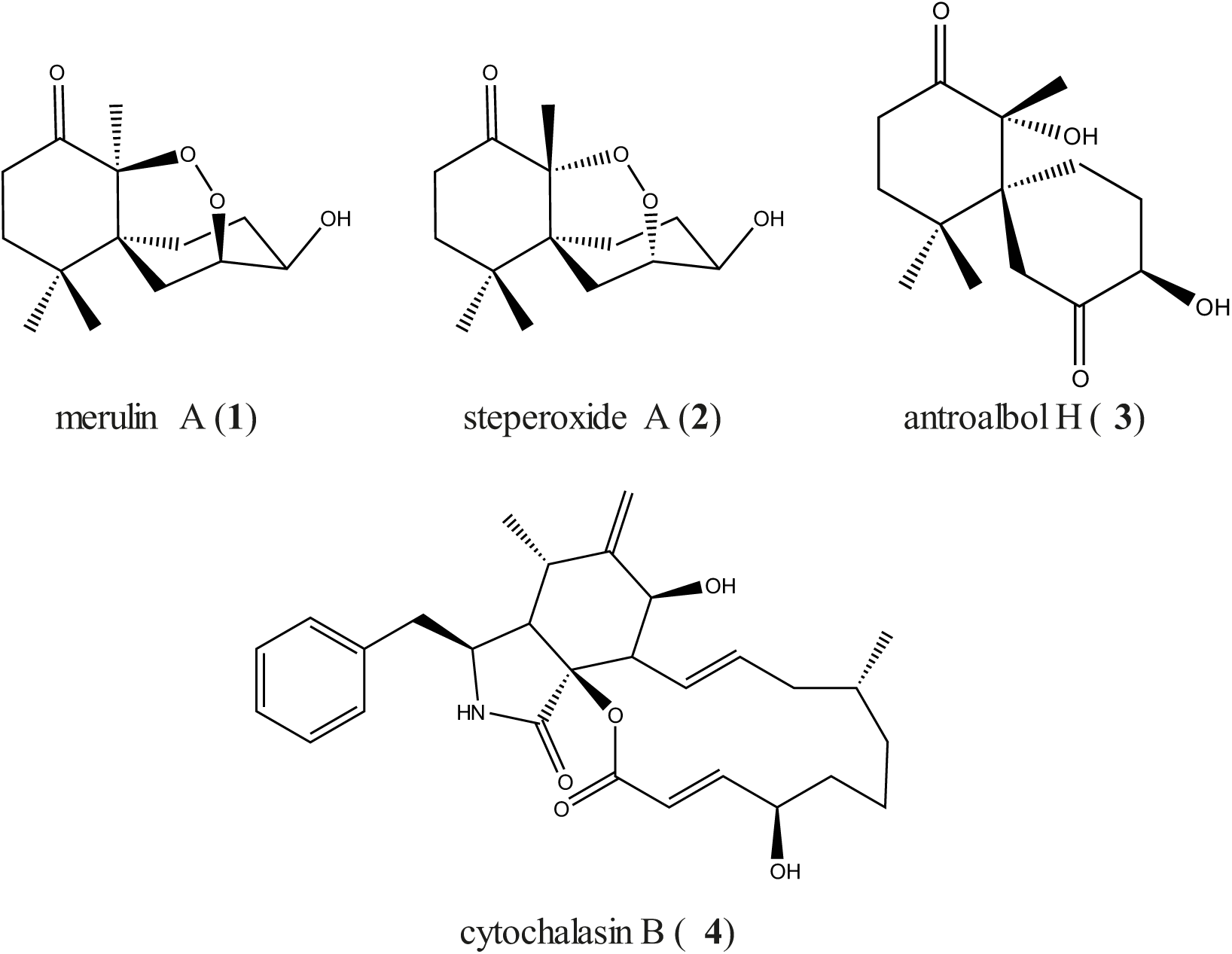
Structures of isolated natural compounds 1-4.

A combination of NMR spectroscopy and mass spectrometry achieved structure elucidation of compounds **1-4**. Compounds **1**-**3**, isolated from *Cerrena zonata* ICMP 16347, were identified as merulin A (**1**) [32], steperoxide A (**2**) [33], and antroalbol H (**3**) [34]. Compound **4**, isolated from *Boeremia sp*. ICMP 17650 was identified as cytochalasin B [35].

Before identification, compounds **1**-**4** were sent to the Community for Antimicrobial Drug Discovery (CO-ADD) at The University of Queensland (Australia) to be evaluated for their antimicrobial activity against a panel of bacterial and fungal pathogens: *Acinetobacter baumannii*, *E. coli*, *Klebsiella pneumoniae*, methicillin-resistant *Staphylococcus aureus* (MRSA), *Pseudomonas aeruginosa*, *Candida albicans* and *Cryptococcus neoformans*. None of the compounds exhibited activity when tested at a concentration of 32 µg/mL (Table 4).

**Table 4.** Antibacterial and antifungal activities of isolated natural compounds as determined by CO-ADD.

| Natural product | Per cent inhibition at 32 µg/mL<sup>a</sup> |  |  |  |  |  |  |
| --- | --- | --- | --- | --- | --- | --- | --- |
|  | *Staphylococcus aureus* | *Escherichia coli* | *Klebsiella pneumoniae* | *Pseudomonas aeruginosa* | *Acinetobacter baumannii* | *Candida albicans* | *Cryptococcus neoformans* |
| Antroalbol H | 12.46 | 4.53 | 11.23 | 16.55 | 10.31 | 1.76 | 26.11 |
| Cytochalasin B | -8.45 | 0.04 | 1.45 | 6.71 | 8.58 | 3.51 | 24.68 |
| Merulin A | 20.57 | 9.56 | 13.69 | 28.92 | 13.16 | 10.13 | 0.51 |
| Steperoxide A | 20.85 | 23.06 | 15.99 | 25.92 | 20.07 | 9.77 | -19.26 |

Key: <sup>a</sup>Data are presented as percentage inhibition (mean of two experiments). *S. aureus* ATCC 43300 (MRSA) with vancomycin (MIC 0.7 µM) as a positive control; *E. coli* ATCC 25922 with colistin (MIC 0.1 µM) as a positive control; *K. pneumoniae* ATCC 700603, *P. aeruginosa* ATCC 27853, and *A. baumannii* ATCC 19606 with colistin (MIC 0.2 µM) as a positive control; *Candida albicans* ATCC 90028 with fluconazole (MIC 0.4 µM) as a positive control; *Cryptococcus neoformans* ATCC 208821 with fluconazole (MIC 26 µM) as a positive control.

## Discussion

In this study, we took a ‘one strain-many compounds’ (OSMAC) approach [18] to screen 32 fungi collected from locations across Aotearoa New Zealand, for activity against *E. coli*. Our pipeline involved growing the fungi on seven media and testing for anti-*E. coli* activity at different ages. Our rationale for taking this approach is the abundant evidence that nutrient sources influence fungal growth and behaviour, including the production of secondary metabolites [18, 19, 36–38].

For our experiments, we grew fungi on media with different pH and various carbon and nitrogen sources (Table 5). These ranged from the defined (Czapek Solution Agar) to the minimal (Water Agar) and included several complex but less chemically defined media made from foods such as oatmeal, potatoes, and rice. Our data provides further strong evidence for the impact of media on both fungal growth and bioactivity. One interesting observation from our data is that some fungi grew faster on water agar than on Czapek Solution Agar and, at times, Czapek Yeast Extract Agar. We consider Water Agar as a minimal nutrient control, as it is only made up of agarose and agaropectin – though there will also be some trace nutrients carried over from the small potato dextrose agar plugs used as inocula. We determined that these small plugs did not affect the pH of the media they were placed on, including water agar. This suggests that Czapek agar-containing media may suppress the growth of some fungi. These were the only media tested that used sodium nitrate as a nitrogen source, so this may be the reason. We could discern no patterns in the fungi affected as they belonged to all three different Phyla, with the two most impacted isolates being the saprophytic Ascomycete *Lauriomyces bellulus* ICMP 15050 and the polypore Basidiomycete *Fomitopsis maire* ICMP 16416.

**Table 5.**
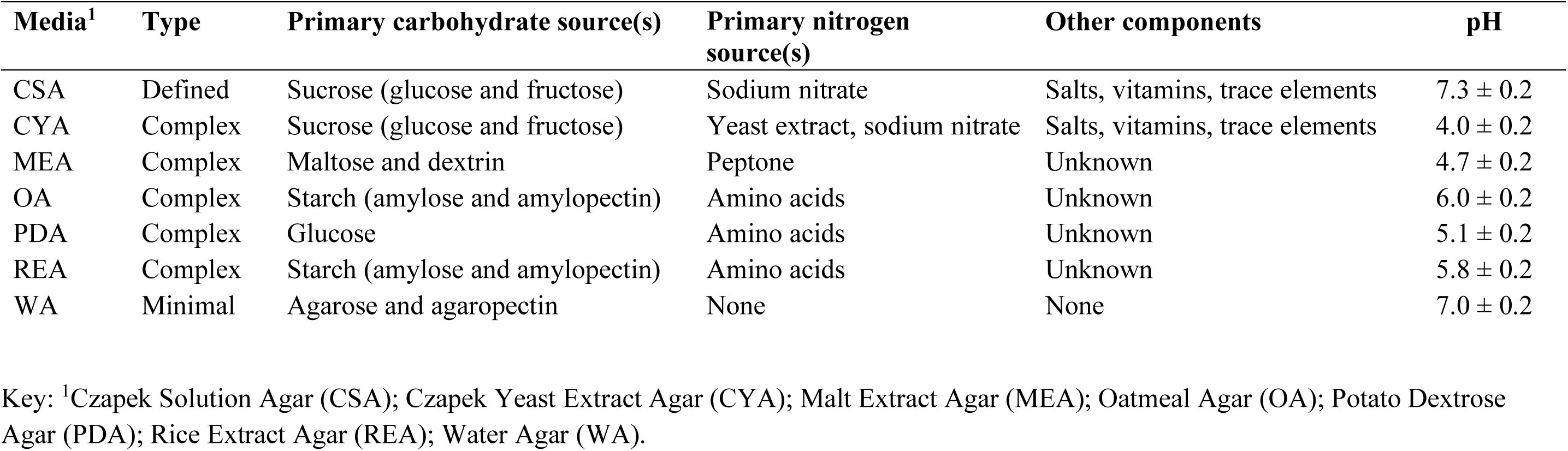
Characteristics of the media used in this study.

Regarding bioactivity, many fungal isolates were active when grown on more than one medium. So far, we have not yet identified any of the compounds responsible for the anti-*E. coli* activity we observed, so whether these fungi are producing the same bioactive compounds in multiple media is an open question. Fungi possess an array of enzymes that enable them to degrade different organic materials, including cellulose and lignin [39, 40]. This likely provides some redundancy in the biosynthetic pathways required for secondary metabolite synthesis. However, as previously described, abundant evidence shows that fungi can produce different secondary metabolites depending on the available nutrients. For example, we found that fungi grown on oatmeal media made more fatty acids, including oleic, palmitic, linoleic, and linolenic acids, as quantified by the intensity of NMR signals. In contrast, these same signals were downregulated by growth on malt extract agar (data not shown).

Similar to our results screening ICMP isolates for anti-mycobacterial activity [17], growth on Czapek Solution Agar resulted in the least anti-*E. coli* activity, while growth on Oatmeal and Potato Dextrose agars resulted in the most activity. However, our results caution against restricting future experiments to Oatmeal and Potato Dextrose agars. If we were to do so, we would likely miss some highly active fungal isolates. For example, *Cunninghamella echinulata* ICMP 1083 produced some of the largest zones of inhibition we observed during this study, but only when grown on Czapek Yeast Extract Agar.

The ICMP fungi we screened in this project included species of several Genera known to produce antimicrobial compounds. For example, Kakoti and colleagues recently isolated cinnabarinic acid from extracts of *Trametes coccinea* mushrooms collected in India [41]. They further showed that cinnabarinic acid could inhibit biofilm formation in *Bacillus subtilis* and *B. cereus*. Chemical extraction and fractionation of a subset of the ICMP isolates, including *T. coccinea*, revealed that much of our observed activity was lost during extraction. However, we identified extracts from *Cerrena zonata*, an unknown species of *Boeremia*, and an unknown species of *Hyaloscypha* (formerly identified as *Pseudaegerita viridis*) that retained activity. We were able to isolate three known compounds from *C. zonata* ICMP 16347 (merulin A [32], steperoxide A [33], and antroalbol H [34]) and one from *Boeremia* sp. ICMP 17650 (cytochalasin B [35]), though none were antibacterial against various human pathogens, including *E. coli*. Investigations are ongoing to identify the source of the anti-*E. coli* activity of *Hyaloscypha* sp. ICMP 16864 and to determine whether this may be due to the production of novel bioactive compounds.

## Author Contributions

BSW, BRC, and SW contributed to the conception and design of the study. SVDP, MMC, ABJG, JMF, and DP performed the experiments. SVDP, MMC, TL, BSW, and SW were involved in data analysis. SVDP and SW wrote the manuscript. All authors contributed to the manuscript revision and read and approved the submitted version.

## Data Availability Statement

The datasets presented in this study can be found in GenBank as detailed in Table 1 and on Figshare (https://auckland.figshare.com/; doi: 10.17608/k6.auckland.28801247 and doi: 10.17608/k6.auckland.28801265)

